# Chlamydiae as symbionts of photosynthetic dinoflagellates

**DOI:** 10.1101/2023.12.18.572253

**Authors:** Justin Maire, Astrid Collingro, Kshitij Tandon, Vanta J. Jameson, Louise M. Judd, Matthias Horn, Linda L. Blackall, Madeleine J. H. van Oppen

## Abstract

Chlamydiae are ubiquitous intracellular bacteria and infect a wide diversity of eukaryotes, including mammals. However, chlamydiae have never been reported to infect photosynthetic organisms. Here, we describe a novel chlamydial genus and species, *Candidatus* Algichlamydia australiensis (*A. australiensis* thereafter), capable of infecting the photosynthetic dinoflagellate *Cladocopium* sp. (originally isolated from a scleractinian coral). *A. australiensis* was confirmed to be intracellular by fluorescence *in situ* hybridization and confocal laser scanning microscopy, and temporally stable at the population level by monitoring its relative abundance across four weeks of host growth. Using a combination of short- and long-read sequencing, we recovered a high-quality (completeness 91.73% and contamination 0.27%) metagenome-assembled genome of *A. australiensis*. Phylogenetic analyses show that this chlamydial taxon represents a new genus and species within the Simkaniaceae family. *A. australiensis* possesses all the hallmark genes for chlamydiae-host interactions, including a complete type III secretion system. In addition, a type IV secretion system is encoded on a plasmid and has previously been observed for only three other chlamydial species. Twenty orthologous groups of genes are unique to *A. australiensis*, one of which is structurally similar to a protein known from Cyanobacteria and Archaeplastida involved in thylakoid biogenesis and maintenance, hinting at potential chlamydiae interactions with the chloroplasts of *Cladocopium* cells. Despite *Cladocopium* being itself a symbiont of cnidarians, a meta-analysis of 12,009 cnidarian 16S rRNA gene metabarcoding samples only returned five samples with *A. australiensis* sequences, suggesting *A. australiensis* does not associate with cnidarians. Our study shows that chlamydiae infect dinoflagellate symbionts of cnidarians, the first photosynthetic organism reported to harbor chlamydiae, thereby expanding the breadth of chlamydial hosts and providing a new contribution to the discussion around the role of chlamydiae in the establishment of the primary plastid.

## Introduction

The Chlamydiota (also known as chlamydiae) is a phylum of obligate intracellular bacteria infecting eukaryotes [1,2]. Despite their diversity, all known chlamydiae have a remarkably conserved biology; they are dependent on eukaryotic host cells for growth and survival, alternate between infectious extracellular elementary bodies and intracellular replicative reticulate bodies, and manipulate host cells through a type III secretion system (T3SS) [1]. Because of their intracellular lifestyle, all chlamydiae also possess reduced genomes (1-3 Mb) [3], and are not culturable *ex hospite*, making them notoriously difficult to study. While chlamydiae are best known for infecting mammals, their host range is incredibly diverse and includes arthropods, amphibians, sponges, corals, and protists [2]. Thus far, however, there has been no record of photosynthetic organisms harboring chlamydiae. Intriguingly, genomic data suggest that numerous horizontal gene transfer events between chlamydiae and Archaeplastida (*i.e.*, algae and plants) have occurred, which were speculated to be remnants of ancestral interactions [4]. This led to the hypothesis that chlamydiae facilitated the establishment of ancestral Cyanobacteria as plastids [4,5], though phylogenetic evidence for this hypothesis remains controversial [6].

Photosynthetic dinoflagellates of the Symbiodiniaceae family associate with a wide range of intra- and extracellular bacteria [7–10], some of which are beneficial for Symbiodiniaceae physiology and stress tolerance [11–15]. While they can be free-living, Symbiodiniaceae are widely known as endosymbionts of cnidarians, which they provide with organic carbon through photosynthate transfer [16]. Chlamydiae sequences were recently detected in 16S rRNA gene metabarcoding of several Symbiodiniaceae laboratory cultures [7,10,14]. In a *Cladocopium* sp. culture (SCF049.01 - See supplementary text and Figure S1 regarding its taxonomy), a single chlamydial amplicon sequence variant (ASV) made up more than 60% of the associated bacterial communities [7], but the nature of the association or whether *Cladocopium* is the true chlamydial host was not investigated. By combining metabarcoding, fluorescence microscopy, and genome sequencing and analysis, we prove that *Cladocopium* harbors chlamydial cells and provide the first detailed description of an association between a chlamydial representative and a photosynthetic organism.

## Results and discussion

### Chlamydiae reside inside *Cladocopium* cells

To verify that the previously detected chlamydiae infect *Cladocopium* cells, rather than other protists potentially present in the culture, we conducted 18S rRNA gene metabarcoding. All reads from three replicate culture flasks were assigned to Symbiodiniaceae (Figure S2), confirming that there were no other eukaryotes and thus, no other chlamydial hosts, in the culture. Additionally, fluorescence *in situ* hybridization (FISH) and confocal laser scanning microscopy (CLSM) using a chlamydiae-specific probe clearly showed whole bacterial cells fluorescing inside *Cladocopium* cells, as well as on their cell wall (Figures 1A-B). Chlamydiae are usually encased in large host-derived inclusions inside host cells [17], though can sometimes reside in single-cell inclusions [18] or in the cytoplasm [19]. Here, chlamydiae were observed as isolated cells inside *Cladocopium* cells and it is unknown whether these are encased in a host membrane.

**Figure 1:**
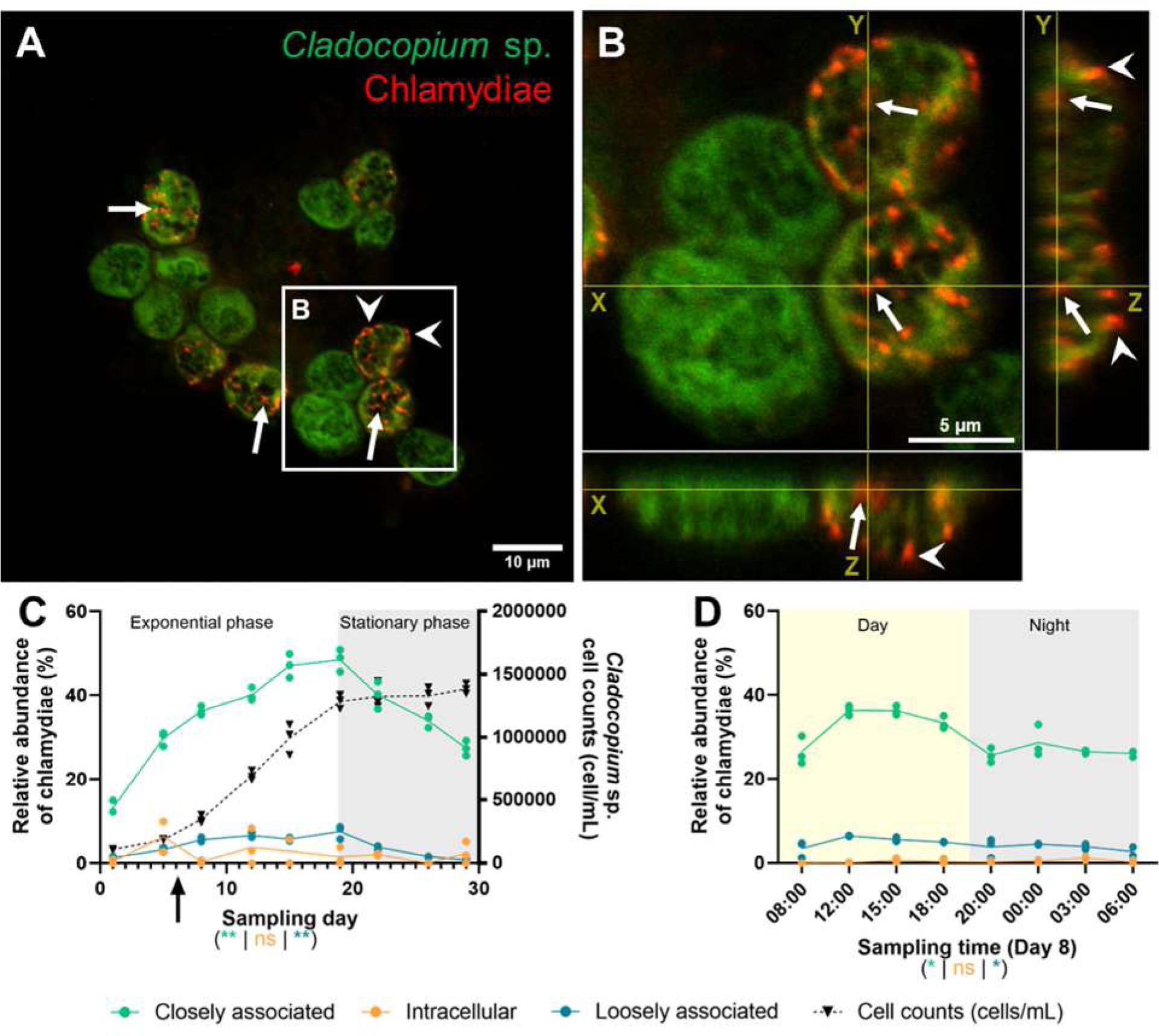
Chlamydiae stably infect *Cladocopium* sp. SCF049.01. **A-B:** Chlamydiae located by FISH inside (white arrows) and on the cell wall (white arrowheads) of *Cladocopium* cells, observed by CLSM. Red: Chls523 probe (chlamydiae); Green: *Cladocopium* autofluorescence. Panel B shows orthogonal projections of a Z-stack of four *Cladocopium* cells from panel A, highlighting the presence of both intracellular and extracellular FISH signal. **C-D:** Relative abundance of chlamydial ASVs in the *Cladocopium* culture across 29 days (C, Growth phase experiment) or 24 h (D, Time series experiment, performed on day 8 of the growth phase experiment - see black arrow in C), determined by 16S rRNA gene metabarcoding (see Figure S4 for experimental design). Bacterial community profiling was performed in three fractions as previously described [7]: loosely associated bacteria (planktonic bacteria; blue), closely associated (intracellular and tightly attached to the cell wall; green) and intracellular bacteria (orange). *Cladocopium* cell counts across the growth phase experiments are also provided (dashed black line, C). Each point represents one of three replicate flasks, with lines connecting the means. For each fraction, the effect of sampling day or time on chlamydial relative abundance is indicated under the plot with its corresponding color, based on a Kruskal-Wallis test. ns: p > 0.05; *: p ≤ 0.05; **: p ≤ 0.01. Single replicate values are available in Table S2.

Absolute quantification of chlamydiae through digital PCR (dPCR) revealed that there were an average of 1716 chlamydial cells/mL of culture supernatant (Figure S3A) and that each *Cladocopium* cell was infected by an average 0.46 chlamydial cells (Figure S3B) suggesting the presence of the common chlamydial developmental cycle with extracellular elementary bodies (EBs) and intracellular replicative bodies (RBs). Because not all *Cladocopium* cells harbored chlamydiae (Figure 1A-B), we measured the number of *Cladocopium* cells stained by FISH (both intracellularly and attached to the cell wall) by flow cytometry across a host growth cycle (once a week for four weeks, Figure S4). This showed that an average of 30% of cells were infected by chlamydiae across four time points (Figure S3C). Therefore, using our previous chlamydial absolute quantification, each infected *Cladocopium* cell (*i.e.,* 30% of the total number of cells) harbors an average 1.52 chlamydial cells (Figure S3B).

### *Cladocopium*-chlamydiae infection is temporally stable

The temporal stability of the association was assessed at the population level by characterizing the bacterial communities throughout a host growth cycle (twice a week for four weeks) and across a single day (eight time points across 24 hours, including four during the day and four at night) (Figure S4, Table S1). Bacterial communities were sampled in three fractions as previously described [7]: loosely associated (planktonic bacteria), closely associated (intracellular and tightly attached to the cell wall), and intracellular. In both experiments, a single chlamydial ASV made up the majority of the reads (Table S2). Chlamydial relative abundance was highest in the closely associated fraction, where it increased from 13% to 48% during host exponential phase and decreased once stationary phase was reached (Figures 1C and S5A, Table S2A). The same trend was observed in the loosely associated fraction, with a peak at 7.5% in relative abundance during host exponential phase. These findings suggest chlamydiae may replicate or be transmitted more easily when their host cells are dividing. The relative abundance of closely associated chlamydiae was higher during the day (33.1% across the four time points) than at night (26.7%) (Figure 1D, S5B, Table S2B). This may be due to the lack of host photosynthesis at night, resulting in lower amounts of ATP available for chlamydial survival and replication. Chlamydial relative abundance in the intracellular fraction was very low (< 5%) in both experiments, suggesting EBs might be more abundant than RBs in this culture.

### *Cladocopium*-associated chlamydiae belong to an undescribed Simkaniaceae genus

Using a combination of long- and short-read sequencing, we recovered a metagenome-assembled genome (MAG) of the *Cladocopium*-associated chlamydiae (Cla049; Table 1). The Cla049 MAG is comprised of 1,799,045 bp across seven contigs and estimated to be 91.73% complete. The MAG also includes one 87,402-bp plasmid (pAa). The last common ancestor of all chlamydiae likely possessed a plasmid, which was lost and/or integrated into the chromosome of some chlamydial lineages, but conserved in others [20,21]. Cla049’s plasmid possesses key genes present on other chlamydial plasmids, including *pgp1* (a replicative DNA helicase) and *pgp2* (a virulence protein) (Table S3). Other genes typically present on chlamydial plasmids, such as *praA*/*pgp5* (a chromosome partitioning protein) and *pgp6* (involved in host cell response mediation), are on non-circular contigs and therefore may have been integrated into the genome. Cla049 encodes 11 transposases, including two on its plasmid, hinting at a potential for gene flow between the plasmid and chromosome that may explain the presence of plasmid genes on non-circular contigs.

**Table 1:** Summary statistics from the *Candidatus* Algichlamydia australiensis genome (Cla049 MAG).

| Genome | Cla049 |
| --- | --- |
| Size (bp) | 1,799,045 |
| Long-read coverage (X) | 10 |
| Short-read coverage (X) | 2,253 |
| Completeness (%) | 91.73 |
| Contamination (%) | 0.27 |
| Estimated complete size (Mb) | 2.0 |
| G+C content (%) | 42.1 |
| N50 | 497,235 |
| Number of contigs | 7 |
| Number of coding sequences (CDSs) | 1,644 |
| Number of ribosomal RNA operons | 1 |
| Number of transfer RNAs | 21 |
| Plasmids | 1 (pAa) |
| Plasmid size (bp) | 87,402 |

To assess the taxonomic placement of the Cla049 MAG, we constructed two maximum likelihood phylogenetic trees based on 15 conserved marker genes (Table S4) and 169 chlamydial genomes (Table S5), calculated with IQ-TREE using different models (Figures 2 and S6). Cla049 represents an undescribed, deep-branching genus within the Simkaniaceae family. Cla049 is most closely related to a group of three MAGs, including two isolated from activated sludge in Hong Kong (HK-STAS-VERR_A-4 and HK-STAS-VERR_A-5), and one from a marine microbial biofilm in Norway (OFTM343). Average amino acid identity (AAI) is ∼40-47% with all Simkaniaceae and Parasimkaniaceae genomes (Table S6), supporting the placement of Cla049 into a novel genus and species, which we propose to name *Candidatus* Algichlamydia australiensis gen. nov. sp. nov. (*A. australiensis* Cla049 hereafter). The name has been registered through SeqCode [22]

**Figure 2:**
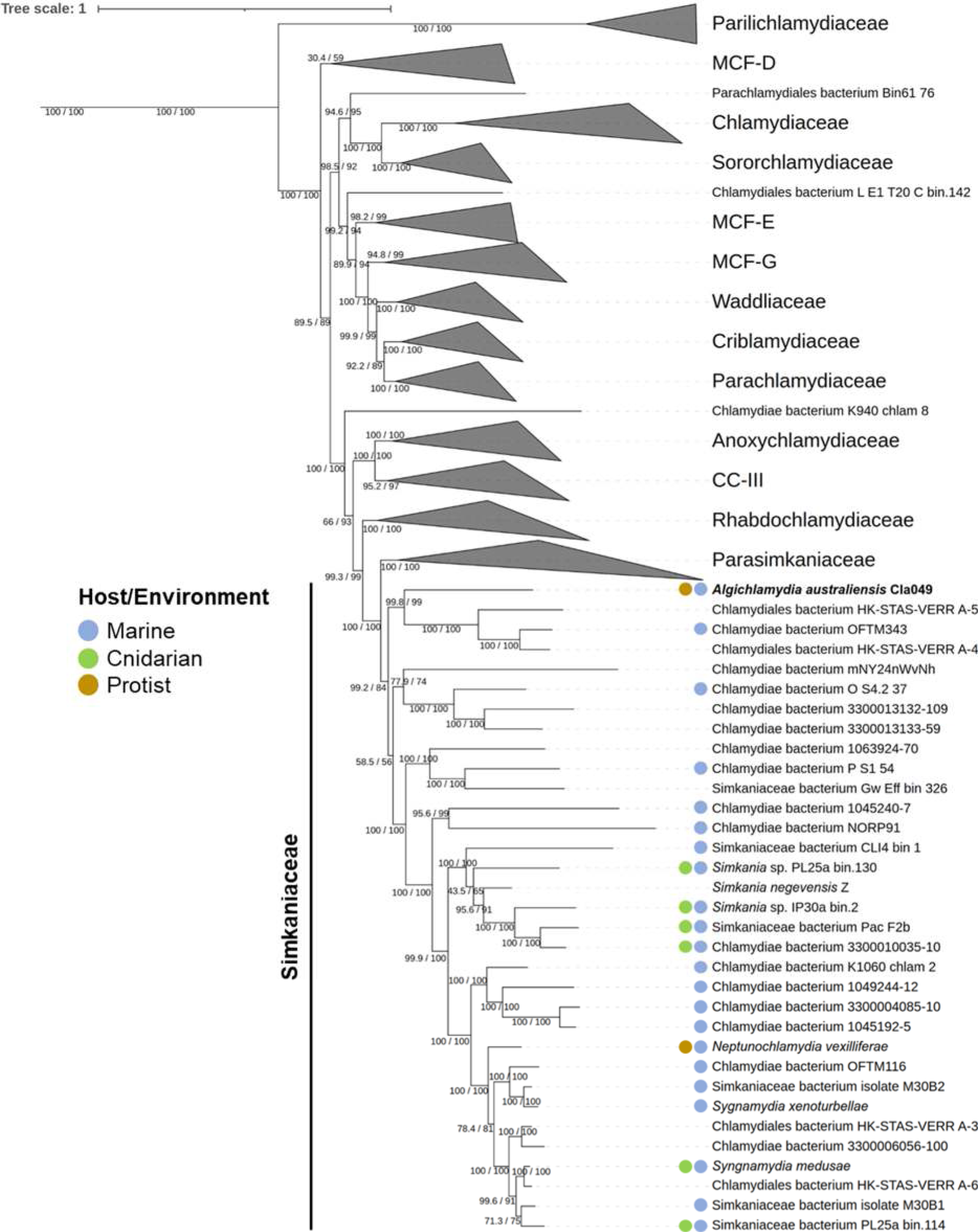
The chlamydial symbiont *Candidatus* Algichlamydia australiensis (Cla049 MAG) belongs to an undescribed, deep-branching Simkaniaceae genus. Chlamydial maximum likelihood phylogeny based on 15 conserved gene markers (Table S4) in 169 chlamydial genomes (Table S5). Confidence values based on 1000 ultrafast bootstrap replicates and 1000 replicates of the SH-like approximate likelihood ratio test are provided. This tree was calculated using IQ-TREE 2 [23] with ModelFinder Plus under the LG+C20+R4 model [24]; an additional tree calculated under the posterior mean **s**ite frequency (PMSF) model [25] using this tree as seed confirmed the phylogeny and is available in Figure S6. MCF: metagenomic chlamydial family; CC-III: chlamydiae clade III.

### *Algichlamydia australiensis* Cla049 has a reduced metabolic potential

The metabolic potential of *A. australiensis* is heavily reduced (Tables S7 and S8), lacking pathways for the synthesis of nucleotides, vitamins, and most amino acids (only genes for glutamate, aspartate, lysine, and alanine biosynthesis were found, as well as glycine-serine interconversion). The *A. australiensis* genome contains the genes necessary for glycolysis, the tricarboxylic acid cycle, pyruvate oxidation, the pentose phosphate pathway, and glycogenesis. *A. australiensis* also possesses a complete shikimate pathway, as well as a complete menaquinone biosynthesis pathway, a cofactor that promotes growth in *Chlamydia trachomatis* [26]. Finally, the genome was predicted to produce two secondary metabolites, most closely related to nostovalerolactone (Table S9), a putative transcriptional regulator [27]. These metabolic abilities, or lack thereof, are similar to other chlamydiae [28,29], and suggest that *A. australiensis* acquires most metabolites and energy from its *Cladocopium* host.

*A. australiensis* was predicted to encode five nucleotide transport proteins (NTTs), which were all most closely related to the four NTTs of *Simkania negevensis*, the Simkaniaceae type species (Figure S7). NTTs are critical for chlamydiae to import essential nutrients, including ATP, from their host cell [30,31]. In *S. negevensis*, *Sn*NTT1 is an ATP/ADP antiporter, *Sn*NTT2 is a guanine nucleotide/ATP/H^+^ symporter, *Sn*NTT3 acts as an RNA nucleotide antiporter, and no substrate was identified for *Sn*NTT4 [31]. However, *Sn*NTT1’s putative ortholog in *A. australiensis* Cla049 was split into three CDSs (EFCAOE_00685/00690/00695), with a STOP codon at the end of EFCAOE_00690 and a STOP codon and a frameshift at the end of EFCAOE_00695, and is thus presumably not functional, drastically diminishing the potential for ATP import and consequently energy parasitism. Nonetheless, *A. australiensis* Cla049 appears to possess two orthologs of *Sn*NTT3 (Figure S7; EFCAOE_00240 and EFCAOE_06465), which can import ATP in exchange for other nucleotides and may therefore compensate for the hypothetical non-functionality of NTT1. Functional analyses are needed to determine the true substrates of *A. australiensis* Cla049’s NTTs. Alternatively, because *Cladocopium* sp. is photosynthetic, it may have lower intracellular ATP and higher intracellular glucose than heterotrophic host cells, and *A. australiensis* Cla049 may rely more on glycolysis than ATP parasitism.

### The chromosome and plasmid of *Algichlamydia australiensis* Cla049 encode a T3SS and type IV secretion system (T4SS), respectively

*A. australiensis* Cla049 encodes most chlamydial hallmark virulence-associated genes that are present in other chlamydiae (Table S10) [28]. This includes adhesins, T3SS effectors, Ser/Thr kinases (for host cell modulation), and developmental regulators. We also found 17 genes encoding putative eukaryotic-like repeat proteins (12 ankyrin repeat proteins, one WD40 repeat protein, and four tetratricopeptide repeat proteins), which are putative mediators of host-bacteria interactions and also abundant in other chlamydiae [21]. Finally, *A. australiensis* Cla049 encodes a complete T3SS and T4SS (Figure 3), the latter being encoded on the plasmid. Both secretion systems show high synteny with the corresponding coding regions of the chromosome and plasmid, respectively, of *S. negevensis* (Figure 3). The chlamydial T3SS is highly conserved and allows for the translocation of effectors into eukaryotic host cells [32]. The T4SS has only been reported in Simkaniaceae and Parachlamydiaceae, and its role remains unclear, though it may be involved in plasmid propagation [20,21,33]. The chlamydial T4SS is plasmid-encoded in only three other known chlamydial species, *S. negevensis*, *Protochlamydia naegleriophila*, and *Rubidus massiliensis*, while T4SS-encoding genes have been integrated in the chromosome of many Parachlamydiaceae and Simkaniaceae species [20,21]. It was hypothesized that a T4SS of alphaproteobacterial origin was integrated into the plasmid of the Parachlamydiaceae ancestor and subsequently acquired by the plasmid of the Simkaniaceae ancestor [20]. The presence of a plasmid-encoded T4SS in two distantly related Simkaniaceae species, *S. negevensis* and *A. australiensis* Cla049, supports this hypothesis.

**Figure 3:**
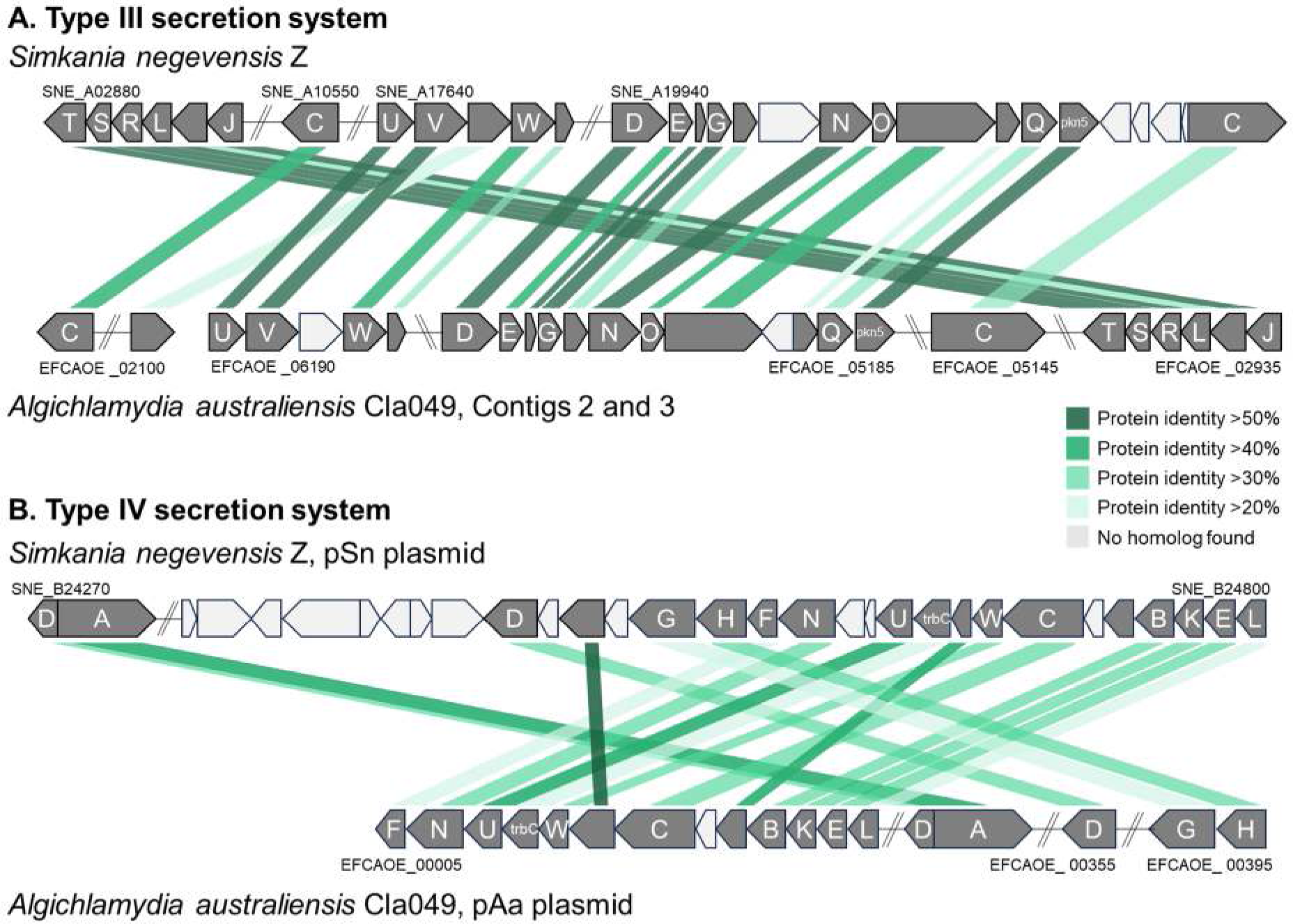
Completeness, similarity and synteny of the genomic regions encoding the T3SS (A, *sct* genes) and T4SS (B, *tra* genes) in *Algichlamydia australiensis* Cla049, compared with *Simkania negevensis* Z. Synteny is shown by lines joining the two genomes; protein identity is color-coded.

### Unique genes in *A. australiensis* Cla049 may impact host-symbiont interactions

Because *A. australiensis* Cla049 is the first confirmed chlamydial symbiont of a photosynthetic organism, we asked whether it possessed any unique genes, not present in other known chlamydiae, which may mediate interactions with a photosynthetic host. We found 20 orthologous groups (OGs) that were specific to *A. australiensis* Cla049 (Table S11). Only one OG (OG0007644) was reliably annotated and was predicted to encode a lanthionine synthetase. Lanthionine is a major component of lantibiotics, a class of antimicrobial compounds [34]. Some chlamydiae defend their hosts against pathogens; for example *Parachlamydia acanthamoebae* protects its amoeba host against giant viruses [35]. Symbiodiniaceae harbor a wide range of viruses, including filamentous [36] and megaviruses [37]. Comparisons of viral diversity and abundance between *Cladocopium* cultures cured of *A. australiensis* Cla049 (*e.g.,* through rifampicin treatment) and control cultures may shed light on any potential defensive role through, for example, lantibiotic production.

Other OGs were predicted to encode hypothetical proteins without any known motifs. Thus, we predicted the structure of one representative sequence per OG using AlphaFold and queried the RCSB Protein Data Bank to identify structurally similar proteins. Only four genes resulted in reliable structural predictions (pLDDT > 70; Table S11 and Figure S8). Among them, one gene (EFCAOE_07285) had two particularly relevant close hits: (i) a putative ankyrin repeat protein, which may be involved in host-chlamydiae interactions; (ii) a vesicle-inducing protein in plastids 1 (VIPP1), a protein involved in the biogenesis and integrity of thylakoid membranes of plants, algae, and cyanobacteria [38]. In *Arabidopsis thaliana*, VIPP1 knockdown results in thylakoid swelling under high light [39], and VIPP1 overexpression improves post-heat stress recovery [40]. Therefore, EFCAOE_07285 may mediate interactions between *A. australiensis* Cla049 and *Cladocopium*’s thylakoids, and impact the photophysiology of *Cladocopium* cells.

### *In hospite* Symbiodiniaceae may not harbor *Algichlamydia australiensis* Cla049

Our previous study of Symbiodiniaceae-associated bacteria showed that ASVs at least 99% identical to *A. australiensis* Cla049 also infects cultures of *Gerakladium* sp. G3, *Breviolum minutum*, *Durusdinium trenchii*, and *Fugacium* sp. F5.1, though at much lower abundances than in *Cladocopium* sp. SCF049.01 (Table S12A) [7]. Because these Symbiodiniaceae cultures are themselves symbionts of cnidarians, we asked whether *A. australiensis* Cla049 infects *in hospite* Symbiodiniaceae (within cnidarians), or cnidarians themselves that are known to be chlamydial hosts [2,41–43]. Analysis of a recent dataset of 12,009 cnidarian 16S rRNA gene metabarcoding samples from 186 studies [44] showed that only five samples (each from a different study) harbored ASVs at least 99% identical to the 16S rRNA sequence of *A. australiensis* Cla049 (Table S12B). The five samples were from four different genera of scleractinian corals, and the relative abundance of *A. australiensis* Cla049 varied between 0.01% and 0.31% (Table S12B). This low abundance and prevalence suggest that *A. australiensis* Cla049 is not a common symbiont of *in hospite* Symbiodiniaceae or cnidarians. Nonetheless, this may be underestimated because of known biases from common universal primers against chlamydiae (*e.g.,* 515F/806R of the V4 region, or 1391R of the V8 region [45]), as well the low numbers of *A. australiensis* Cla049 in *Cladocopium* cells, which may be missed if the sequencing depth is too low.

## Conclusion

We provided a genomic characterization of a member of the chlamydiae that infects cultures of the dinoflagellate *Cladocopium* sp. SCF049.01. As far as we know, this is the first description of a chlamydial infection in a photosynthetic organism [2]. Nonetheless, chlamydial 16S rRNA gene reads were recently detected in cultures of the green alga *Ostreobium* sp. [46,47], and chlamydial MAGs were retrieved from cultures of the green alga *Amoebophrya* sp. and the kelp *Saccharina japonica* [48]. However, in all these examples, it was not established whether the photosynthetic organism or associated protists were the chlamydial host. Thus, additional studies are required to fully appreciate the potential breadth of photosynthetic hosts of chlamydiae. We showed that *A. australiensis* Cla049 is intracellular, temporally stable, belongs to the Simkaniaceae family, and has a similar metabolic and host interaction potential to other chlamydiae. It is only the fourth chlamydial species to possess a plasmid-encoded T4SS, thereby increasing our understanding of the evolution of chlamydial plasmids. Twenty OGs in the *A. australiensis* Cla049 MAG were not found in any other chlamydiae, which included genes putatively involved in the production of antimicrobial compounds or in interactions with host thylakoids. Therefore, our study reveals that chlamydiae can infect photosynthetic hosts. This adds to the enormous diversity of eukaryotic hosts that harbor chlamydiae and provides valuable insight into the evolutionary history of chlamydiae. This first example of a chlamydia infecting a photosynthetic host cell also sheds new light on a hypothesis about a possible contribution of chlamydiae to the establishment of cyanobacterial symbionts during the evolution of the first photosynthetic eukaryotes [4–6].

## Material and methods

### Symbiodiniaceae culture and maintenance

The Symbiodiniaceae culture SCF049.01 was used in this study (see Supplementary text for taxonomic considerations). It was initially isolated at the Australian Institute of Marine Science from the coral *Pocillopora damicornis* collected from Davies Reef (central Great Barrier Reef, Australia). Symbiodiniaceae cultures were maintained in 15 mL Daigo’s IMK medium (1×), prepared with filtered red sea salt water (fRSSW, 34 ppt salinity) in sterile 50 mL polypropylene culture flasks where media was changed fortnightly. These flasks were kept in a 12 h light:12 h dark incubator (50–60 μmol photons m^−2^ s^−1^ of photosynthetically active radiation) at 26°C.

### FISH on Symbiodiniaceae cells

Symbiodiniaceae cells were fixed in ice-cold 66% ethanol and photobleached as previously described [7]. Teflon-printed microscope slides (ProSciTech) were coated with poly-L-lysine solution (0.01%) in PBS by aliquoting the solution onto printed wells, incubated at 37°C for 3 h, then thrice washed in sterile Milli-Q water and air dried. Sample aliquots of 5-10 μL were pipetted into a well on treated ten-well slide and FISH was then performed as previously described [7]. Hybridization was in 16 μL buffer (0.9M NaCl, 20 mM Tris-HCL pH 7.2, 25% formamide, 0.01% SDS) and 2 μL of the 16S rRNA-targeting, chlamydiae-specific probe Chls523 (CCTCCGTATTACCGCAGC; [49]), conjugated with a DOPE-Cy3 fluorophore, along with 2 µL of a competitor probe (CCTCCGTATTACCGCGGC; [49]), both at a final concentration of 5 ng/μL. NaCl concentration in washing buffer was 0.149 M. Slides were mounted with CitiFluor™ CFM3 mounting medium (Hatfield, PA, USA) and stored in the dark at −20 °C until observation by CLSM.

### Confocal Laser Scanning Microscopy (CLSM)

Observations were made on a Nikon AIR CLSM (Nikon, Tokyo, Japan) with the NIS325 Element software. Virtual band mode was used to acquire variable emission bandwidth to tailor acquisition for specific fluorophores. An oil immersion objective of 60X magnification and 1.4 numerical aperture was used for image acquisition. The fluorophore Cy3 was excited using the 561 nm laser line, and the Symbiodiniaceae autofluorescence using the 488 nm laser line, with a detection range of 570-635 nm for Cy3, and 670-720 nm for Symbiodiniaceae. Z-stacks were acquired using Z steps of 0.2 μM. Nd2 files were processed using ImageJ. Single photos were extracted from Z-stacks. Each photo was selected in the center of the Z-stack and covers 0.2 µm. As Symbiodiniaceae cells have a diameter of around 8-10 µm, FISH signal present within a Symbiodiniaceae cell represents intracellular bacteria.

### Determination of chlamydial copy numbers by digital PCR

To determine chlamydial cell numbers present outside (*i.e.,* in the supernatant) of the host and intracellular in *Cladocopium* sp., we performed digital PCR (dPCR). Six separate culture flasks containing 9 mL of cultures were harvested by shaking and subsequently centrifuged at 5,000 × *g* for 5 min. Each sample was then separated in supernatant (n = 6) and cellular fraction (n = 6). The supernatant was filtered through 5 µm and 1.2 µm filters to retrieve extracellular chlamydiae only. *Cladocopium* sp. cells in the cellular fraction were washed once in IMK medium and subsequently counted with a LUNA-FX7 cell counter (Logos Biosystems). DNA was extracted from all samples using the DNeasy PowerSoil Pro Kit (Qiagen) according to the manufacturer’s instructions. Only the initial cell lysis step was carried out in a FastPrep-24 instrument in lysing matrix A tubes (both MP Biomedicals) under different settings for the supernatant and cellular fraction samples, respectively (supernatant: 4.0 m/sec, 30 seconds; cellular fraction: 5.0 m/sec, 40 seconds). Resulting DNA concentrations were determined with Nanodrop and diluted to roughly use 0.5 ng/µL DNA as template for the dPCR. Digital PCR was performed using the QIAcuity EvaGreen PCR kit (Qiagen) and the chlamydiae-specific primer pair Chl40F (5’ CRG CGT GGA TGA GGC AT 3’) and Chl523R (5’ CCY YMC GTA TTA CCG CAG CT 3’) targeting the 16S rRNA gene on a QIAcuity One digital PCR device (Qiagen) as recommended by the manufacturer. The cycling conditions were 95°C for 2 min; 35 cycles of 95°C for 30 sec, 64.5°C for 45 sec, 72°C for 1 min; 40°C for 5 min with imaging at an exposure time of 350 ms and a gain of 3. Direct quantification data were acquired with the QIAcuity Software Suite 2.2.0.26 (Qiagen) and chlamydial 16S rRNA copy numbers per mL were calculated based on these data. As the *A. australiensis* Cla049 genome contains only one copy of the 16S rRNA gene, the determined gene copy numbers per mL correspond to chlamydial cell numbers per mL. In the cellular fraction, the chlamydial cell numbers per mL were normalized to the *Cladocopium* sp. cell counts.

### Growth phase and time series experiments (for metabarcoding and flow cytometry)

To analyze *Cladocopium*-associated bacterial communities through time, the *Cladocopium* sp. culture was sub-cultured into three separate flasks at 1 × 10^5^ cells/mL in 100 mL 1X IMK media on day 1. For the ‘growth phase’ experiment, all three flasks were sampled on days 1, 5, 8, 12, 15, 19, 22, 26, 29, until the culture reached stationary phase (Figure S4). All sampling was done between 14:00 and 16:00. On day 8-9, during exponential phase, all three flasks were sampled across eight time points across the two days for the ‘time series’ experiment (Figure S4): 08:00 (day 8), 12:00, 15:00, 18:00, 20:00, 00:00 (day 9), 03:00, 06:00. Samples for day 8 of the growth phase experiment and for the 15:00 timepoint of the time series experiment were the same.

For each sampling day/time, cell density was obtained using a Countess II FL cell counter (Thermo Fisher scientific). Samples (100 μL) were taken from each of the three replicate flasks. Of this, two 10 μL samples were processed to obtain cell density for each flask. Cultures were then sampled as previously described [7] in order to separate “loosely associated”, “closely associated”, and “intracellular” bacteria from *Cladocopium* sp SCF049.01 cultures. Briefly, six replicates of 100,000 cells for each flask were filtered through a 5-µm strainer (pluriSelect, Germany), which retains *Cladocopium* cells (> 5 µm), but not planktonic bacteria (< 5 µm). Three replicates were washed with fRSSW to wash away bacteria that are not tightly attached to the cell surface. Filtrates represent the “loosely associated bacteria”. Filters were detached from the strainers and represent the “closely associated bacteria”, *i.e.* intracellular bacteria and those tightly attached to the surface. The other three replicates were washed with 6% sodium hypochlorite (v/v) to wash away all extracellular bacteria. Filters were detached from the strainers and these represent the “intracellular bacteria”. Six filters (three for the growth phase experiment, three for the time series experiment) that only received fRSSW and six filters (three for the growth phase experiment, three for the time series experiment) that only received sodium hypochlorite (no algae) were also sampled as negative controls. All samples were snap-frozen and kept at - 20°C until processing for metabarcoding.

Additional samples were taken during the growth phase experiment for FISH and flow cytometry quantification of *Cladocopium* cells infected by chlamydiae, at days 1, 8, 15, 22, and 29. Approximately 3 ×10^6^ *Cladocopium* cells from each flask were sampled for each time point and fixed in 80% ethanol as previously described [7].

### FISH and flow cytometry

FISH and flow cytometry analyses were conducted as previously described [7]. Each sample was separated into two aliquots. FISH was performed on both aliquots directly in the tube, one receiving no probe during the hybridization step, while the other received the 16S rRNA-targeting, chlamydiae-specific probe Chls523 (CCTCCGTATTACCGCAGC; [49]), conjugated with a DOPE-Cy3 fluorophore, along with a competitor probe (CCTCCGTATTACCGCGGC; [49]), both at a final concentration of 5 ng/μL. Samples were analyzed on a CytoFLEX LX flow cytometer (Beckman Coulter Inc, USA). For each sample, the unstained aliquot was used to determine cell autofluorescence in the FISH-specific channel (561 nm laser; emission 610 ± 10 nm), and a quadrant was drawn to encompass unstained cells on one side (autofluorescence only), and stained cells on the other side (autofluorescence + FISH signal) [7]. The proportion of cells above the line was interpreted as the proportion of cells stained by FISH.

### DNA extractions for metabarcoding

DNA extractions were performed using a salting-out method with modifications as previously described [50]. Extraction blanks (two to four) were included to account for potential contaminants introduced during the extraction process.

### Metabarcoding library preparation

For Symbiodiniaceae profiling, the ITS2 region was amplified using the primer pair Sym_Var_5.8S2 (5’ <u>GTGACCTATGAACTCAGGAGTC</u>GAATTGCAGAACTCCGTGAACC 3′) and Sym_Var_Rev (5’ <u>CTGAGACTTGCACATCGCAGC</u>CGGGTTCWCTTGTYTGACTTCATGC 3′) [51]. For 18S rRNA profiling, the primer pair 574*F (<u>GTGACCTATGAACTCAGGAGTC</u>CGGTAAYTCCAGCTCYV) [52] and 952R (<u>CTGAGACTTGCACATCGCAGC</u>TTGGCAAATGCTTTCGC) [53] was used. For bacterial communities, hypervariable regions V5-V6 of the 16S rRNA genes were amplified using the primer pair 784F (5ʹ <u>GTGACCTATGAACTCAGGAGTC</u>AGGATTAGATACCCTGGTA 3ʹ) and 1061R (5ʹ <u>CTGAGACTTGCACATCGCAGC</u>CRRCACGAGCTGACGAC 3ʹ). Underlined are the Illumina adapters attached to the primers. PCR amplification, library preparation, and sequencing were completed as previously described [54]. Four no-template PCRs were included per primer pair. Sequencing was performed on an Illumina MiSeq platform using v3 (2 × 300 bp) reagents at the Walter and Eliza Hall Institute (Melbourne, Australia).

### Metabarcoding data analyses

Symbiodiniaceae metabarcoding data (ITS2) were processed using the SymPortal analytical framework (symportal.org) [55]. Sequence information was submitted to the SymPortal remote database and underwent quality control including the removal of artefact and non-Symbiodiniaceae sequences. Relative abundances of ITS2 types were exported and plotted on GraphPad Prism 9.

18S rRNA gene metabarcoding data were processed using QIIME2 version 2021.8 [56]. Metabarcoding data were obtained as paired-end, demultiplexed files with primers and adapters attached. The cutadapt plugin [57] was used to remove primer and adapter sequences, with an error rate of 0.2. The quality of trimmed sequences was determined using the DADA2 plugin [58], which denoises, filters, dereplicates, detects chimeras and merges paired-end reads. Reads with low quality (Q-score < 30) were removed. Taxonomy was assigned by training a naive Bayes classifier with the feature-classifier plugin [56], based on a 99% similarity to the 18S rRNA gene in the PR^2^ database v4.14.1 to match the primer pair used [59]. Metadata file, phylogenetic tree, and tables with Amplicon Sequence Variant (ASV) taxonomic classifications and counts were exported, and relative abundances were plotted using GraphPad Prism 9.

16S rRNA gene metabarcoding data were processed using QIIME2 version 2021.8 [56]. Metabarcoding data were obtained as paired-end, demultiplexed files with primers and adapters attached. The cutadapt plugin [57] was used to remove primer and adapter sequences, with an error rate of 0.2. The quality of trimmed sequences was determined using the DADA2 plugin [58], which denoises, filters, dereplicates, detects chimeras and merges paired-end reads. Reads with low quality (Q-score < 30) were removed. Taxonomy was assigned by training a naive Bayes classifier with the feature-classifier plugin [56], based on a 99% similarity to the V5-V6 region of the 16S rRNA gene in the SILVA 138 database to match the 784F/1061R primer pair used [60]. Mitochondria and chloroplast reads were filtered out. The metadata file, phylogenetic tree, and ASV tables were imported into Rstudio for analyses, using the phyloseq package [61]. At this stage of the analysis, datasets from the growth phase experiment and from the time series experiment were separated and analysed independently. Rare ASVs (percentage abundance lower than 1 × 10^-5^) were removed from the dataset. Samples with low read numbers were removed (four for the growth phase experiment: Day 5 - Intracellular - Flask C, Day 15 - Intracellular - Flask B, Day 22 - Intracellular - Flask A, Day 26 - Intracellular - Flask B; three for the time series experiment: 12:00 - Loosely associated - Flask C, 0:00 - Loosely associated - Flask A, 6:00 - Loosely associated - Flask B). Sequencing statistics for all metabarcoding experiments can be found in Table S1. Contaminant ASVs, arising from kit reagents and sample manipulation, were identified manually based on their abundance in negative controls: any ASV that was five times more abundant in the mean abundance of either filter blanks, extraction blanks or no template PCRs compared to the mean of all *Cladocopium* sp SCF049.01 samples, and that represented at least 500 reads in all *Cladocopium* sp SCF049.01 samples, was considered a contaminant, and removed from the dataset. Known contaminants (*e.g.*, *Cutibacterium*) were also removed manually. For the growth phase experiments, 299 ASVs were identified as contaminants, accounting for 5.99% of *Cladocopium* sp SCF049.01 reads (Table S13A). For the time series experiment, 33 contaminants were identified, accounting for 9.88% of *Cladocopium* sp SCF049.01 reads (Table S13B). ASVs assigned to the chlamydiae were specifically targeted and plotted separately.

### Statistical analysis

Chlamydial abundance and the proportion of infected *Cladocopium* cells in the growth phase and time series experiments were analyzed and plotted using GraphPad Prism 9. Each experiment was analyzed independently. Within each experiment, the three different fractions (closely associated, intracellular, loosely associated) were analyzed independently. For each fraction, the effect of sampling time (day in the growth phase experiment, hour in the time series experiment) on chlamydial abundance and the proportion of infected *Cladocopium* cells was analyzed by performing a non-parametric Kruskal-Wallis test. Statistical tests were considered significant at α = 0.05, unless otherwise stated.

### Sample preparation and DNA extraction for long-read sequencing of *Cladocopium* microbial communities

For long-read sequencing, the supernatants of SCF049.01 cultures were sampled to minimize host DNA quantities. Two weeks after subculturing (*Cladocopium* density was around ∼1 × 10^6^ cells/mL), 150 mL of supernatant were carefully removed, without disturbing the attached *Cladocopium* cells, to minimize *Cladocopium* contamination. The supernatant was subsequently filtered at 5 µm and 1.2 µm to remove *Cladocopium* cells and > 1.2-µm sized bacteria (chlamydiae are < 1 µm). The filtered supernatant was centrifuged for 15 min at 3,000 × *g* to pellet the bacteria. The supernatant was discarded and the bacterial pellet kept at −20°C. The bacterial pellet was lysed (lysozyme 100 mg/mL) and DNA was extracted with GenFind V3, according to manufacturer’s instructions for bacterial samples (Beckman Coulter). A ligation sequencing library was prepared (ONT SQK-NBD114-96) and the resultant library run on a MinION flow cell (FLO-MIN114) using a GridION device. Data was basecalled with Super-accurate basecalling in MinKNOW v23.07.05.

### Sample preparation and DNA extraction for short-read sequencing of *Cladocopium* microbial communities

For short-read sequencing, we employed a more comprehensive strategy to eliminate *Cladocopium* cells and DNA and maximize bacterial DNA yields. First, the ‘closely-associated’ communities of around 5 × 10^6^ *Cladocopium* cells were sampled as described above. Samples were then bead-beaten for 20 min at 30 Hz with 100 mg of sterile beads (400-600 nm) to open up the *Cladocopium* cells and stained with SYBR Green as previously described [11,14]. Bacteria were then separated from *Cladocopium* cells and debris by fluorescence-activated cell sorting on a Aria III/FACS DiVa 9 software (BD Biosciences, Franklin Lakes, NJ) equipped with a 70 μm nozzle and run at 70 psi, as previously described [14]. Selection was based on size and high SYBR Green fluorescence. A total of 3.75 million events were obtained.

DNA was extracted using a HostZERO Microbial DNA Kit (Zymo) according to the manufacturer’s instructions. Final elution volume was 40 µL. A volume of 20 µL of purified DNA was concentrated using a DNA Concentrator Kit (abcam) according to the manufacturer’s instruction, with a final elution volume of 3 µL. The resulting DNA was amplified by multiple displacement amplification (MDA) using a REPLI-g Single Cell Kit (QIAGEN) following the manufacturer’s instructions. Following MDA at 30°C for 8 h, the DNA polymerase was inactivated, and amplified DNA was stored at −20°C. The sample was sequenced across two lanes of a NovaSeq 6000 SP 2 × 150bp flowcell (Illumina, San Diego, CA) at the Ramaciotti Centre for Genomics (UNSW Sydney, Australia).

### Metagenomic analyses

Long-read sequence quality control was performed with NanoPlot v1.41.6 [62]. Raw sequences were trimmed with NanoFilt v2.8.0 [62], with parameters -q 10 -l 300 for quality and read length, respectively. Assembly was performed using Flye v2.9.2 [63] with two approaches, (i) using default genome mode for flye (flye --nano-raw reads) and (ii) metagenome mode (flye –meta –nano-raw reads). Assembly statistics from both approaches were compared and outputs from metagenome mode were used for downstream analysis.

Contig taxonomy was evaluated with CAT v5.3 [64] using default parameters and the NR database, and with GTDB-Tk v2.3.0 using classify_wf [65]. Five contigs affiliated with the Chlamydiota phylum. All contigs were further binned with GraphMB v0.2.5 [66], and three additional contigs were binned along with the five contigs previously identified as chlamydial, resulting in an eight-contig bin.

The bin was subsequently polished using the short-read sequences. Short-read sequences were merged by read direction, and paired-end reads were quality checked using FASTQC v0.11.5 (https://www.bioinformatics.babraham.ac.uk/projects/fastqc/). Reads were trimmed with trimmomatic v0.36 [67] with the following parameters: CROP:145, LEADING:30, HEADCROP:10, MINLEN:120. Short reads were mapped to the bin obtained above using bowtie2 v2.4.2 [68] and Samtools v1.11 [69]. The bin was then polished with the mapped, high-quality short reads using Pilon v1.24 [70], twice successively. A hybrid assembly was attempted but did not result in a bin of higher quality.

Two contigs from the final bin were predicted to be circular. The content of one of the two contigs was not sufficient to function as a plasmid entity and most chlamydiae only have a single plasmid [20], so we attempted to improve the contiguity of the circular contigs and obtain a single plasmid. Short reads were mapped to the two circular contigs using bowtie2 v2.4.2 [68] and Samtools v1.11 [69], and long reads were mapped to the two circular contigs using minimap2 v2.26 [71] and Samtools v1.11 [69]. A hybrid assembly was conducted with the mapped short and long reads and Unicycler v0.5.0 [72], yielding a single plasmid.

Bin quality and taxonomy were assessed using CheckM2 v1.0.2 [73] and GTDB-Tk v2.3.0 [65], respectively. Bin coverage was obtained using CoverM v0.6.1 (https://github.com/wwood/CoverM) using the “genome” option. Average amino acid identity (AAI) of the obtained bin was calculated with FastAAI (https://github.com/cruizperez/FastAAI) against 48 genomes of the Simkaniaceae and Parasimkaniaceae families.

### Phylogenetic analyses

For comparative and phylogenetic analyses, a dataset of high quality chlamydial genomes based on Dharamshi et al. [29] was used. The chlamydial dataset was complemented by additional genomes available on GenBank/ENA/DDBJ on 29.05.2023. In general, only genomes with a completeness > 70%, contamination < 5% (both determined with CheckM2 v1.0.2 [73]), and average nucleotide identities < 95% (determined with FastANI v1.33 [34]) were considered for the final dataset, which included 170 chlamydial genomes and 89 genomes from Planctomycetota and Verrucomicrobiota serving as outgroup (Table S5). To obtain the phylogenetic affiliation of Cla049, a set of 15 concatenated conserved non-supervised orthologous groups (NOGs) was used (Table S4). These 15 NOGs are known to retrieve the same topology for chlamydial phylogeny as the application of larger protein sets [29]. Proteins of the chlamydial and Planctomycetota-Verrucomicrobiota outgroup genomes belonging to the 15 NOGs were aligned with MAFFT v7.520 L-INS-i [74]. The resulting single protein alignments were subsequently trimmed with BMGE v2.0 [75] and concatenated. One chlamydial MAG with low completeness was excluded from phylogenetic analysis (Table S5). Maximum likelihood phylogeny was inferred with IQ-TREE v2.2.5 [23] using ModelFinder Plus (-m MFP -mset LG,LG+C20 -mfreq “ “ -mrate G4,R4) [24] with 1000 ultrafast bootstrap replicates [76] and 1000 replicates of the SH-like approximate likelihood ratio test [77] under the LG+C20+R4 model. The resulting phylogenetic tree was used as guide tree for maximum likelihood tree inference under the posterior site mean frequency (PMSF) model with 100 non-parametric bootstraps [25]. Phylogenetic trees were rooted using the outgroup and visualized with iTOL v6.8.1 [78].

### Genome annotation

Gene prediction was performed in Bakta v1.7.0 [79], KEGG-mapper Reconstruct [80], eggNOG-mapper v2.1.11 [81], and InterProScan v5.55 with Pfam domain annotations [82]. Pfam annotations were used to look for eukaryotic-like proteins (ankyrin-repeat domains, WD40 domains, tetratricopeptide repeat). Secondary metabolites were predicted using antiSMASH v7.0.0 [83]. The presence of specific genes of interest (virulence genes, T3SS, T4SS) were investigated using BLASTp v.2.14.0 [84] and known sequences from the genomes of either *S. negevensis* [21] or *C. trachomatis* [85].

The genome annotation revealed the presence of five nucleotide transport protein (NTT) genes with one of them being a potential pseudogene separated into three CDSs (EFCAOE_00685, EFCAOE_00690, EFCAOE_00695). To decipher the potential transport capacities of this pseudogenized NTT, we merged the three single CDSs into one combined-CDS. All Cla049 NTTs were then added to a previously published dataset of NTTs [86] aligned with MAFFT v7.520 L-INS-i [74], and trimmed with BMGE v2.0 [75]. Maximum likelihood phylogeny of NTTs was inferred with IQ-TREE v2.2.5 [23] using ModelFinder Plus (-m MFP -mset LG,LG+C20 -mfreq “ “ -mrate G4,R4) [24] with 1000 ultrafast bootstrap replicates [76] and 1000 replicates of the SH-like approximate likelihood ratio test [77] under the LG+R4 model. The resulting phylogenetic was midpoint-rooted and visualized with iTOL v6.8.1 [78].

### Analysis of orthologous groups of proteins

For further comparative analysis of the chlamydial genomes, all encoded protein sequences were clustered into orthologous groups (OGs) with OrthoFinder v2.5.5 [87] under default parameters. OGs and their respective eggNOG annotations were merged in RStudio v2023.06.2 and analyzed. OGs present in the bin obtained in this study, but not present in any other chlamydial genome were selected. Because most sequences did not show any similarity to known proteins in public databases, the structures of representative sequences of these OGs were predicted with AlphaFold v2.3.2 [88] and visualized with UCSF ChimeraX v1.6.1 [89]. The resulting best-ranked protein structure models were then searched with structure search against RCSB protein data bank (PDB) [90]. Only hits with global pLDDT > 70 were considered in the results.

### Symbiodiniaceae taxonomy

To assess the taxonomical placement of SCF049.01, DNA was extracted from ∼1 × 10^6^ cells as described in the section ‘DNA extractions for metabarcoding’. Four conserved DNA markers (internal transcribed space region 2 [ITS2], large ribosomal subunit [LSU], partial chloroplast cp23S, mitochondrial cytochrome b [*cob*], and mitochondrial cytochrome oxidase 1 [*cox1*]) were PCR-amplified, sequenced, and concatenated as previously described [91].

The sequences were trimmed and aligned (MAFFT alignment) with other *Cladocopium* spp. concatenated sequences obtained from previous studies [91,92] using Geneious Prime v2019.1.3. The full alignment was stripped of columns containing 99% or more gaps. This alignment was used to generate a maximum likelihood phylogenetic tree with 1000 ultrafast bootstraps using IQ-TREE v2.2.2.3 [23] with the best model HKY+F+G4, selected by ModelFinder wrapped in IQ-TREE [24].

## Acknowledgments

This research was supported by the Australian Research Council (FL180100036 to MJHvO), the Austrian Science Fund FWF (P32112 to AC), the CDF Visiting Fellowship Program by the Australian Institute of Marine Science (to MH and AC), an Early Career Research grant from the University of Melbourne (to JM), a Global Collaboration Award from the University of Melbourne (to JM), and the Native Australian Animals Trust (to JM). KT is supported by the Australian Research Council (DP200101613). We thank: the Melbourne Research Cloud (University of Melbourne) and the Life Science Compute Cluster (University of Vienna) for providing the high-performance computing instances needed for this work; the Biosciences Microscopy Unit (University of Melbourne) and the Biological Optical Microscopy Platform (University of Melbourne) for the use of their confocal microscopes; Dr Ashley Dungan and Sarah Jane Tsang Min Ching for assistance with molecular work.

## Data availability

All data needed to evaluate the conclusions in the paper are present in the paper and/or the Supplementary Materials.

Raw data are available under the following NCBI BioProject IDs: PRJNA935927 (MiSeq raw data for Growth phase and Time series experiments); PRJNA1018284: (MiSeq raw data for 18S rRNA and ITS2 gene metabarcoding of *Cladocopium* sp. SCF049.01); PRJNA1036848 (raw NovaSeq and minION metagenome sequencing, and the Cla049 MAG [JAXCHI000000000]).

Individual sequences are available under the following NCBI GenBank IDs: *Cladocopium* sp. SCF049.01 ITS2 gene: OR564117; *Cladocopium* sp. SCF049.01 LSU gene OR564118; *Cladocopium* sp. SCF049.01 cp23S gene: OR575593; *Cladocopium* sp. SCF049.01 *cob* gene: OR568575; *Cladocopium* sp. SCF049.01 *cox1* gene: OR568576; *Algichlamydia australiensis* Cla049 16S rRNA gene: OR835247.

