## Supplementary material for "Chlamydiae as symbionts of photosynthetic dinoflagellates"

### Supplementary text

#### Taxonomy of *Cladocopium* sp. SCF049.01

The *Cladocopium* genus has recently seen multiple taxonomic updates, including the formal description of four new species since 2021 [1,2]. Therefore, we assessed whether the culture used in the present study, SCF049.01, fell within any of these newly described species. Phylogeny based on five genetic markers, ITS2, LSU, cp23S, *cox1*, and *cob* placed SCF049.01 within the C1 radiation; it is most closely related to the newly described *C. proliferum* and *C. vulgare* (Figure S1A). Metabarcoding of the ITS2 region confirmed its ITS2 type as C1 (Figure S1B). Surprisingly, it is more distantly related to *C. latusorum* and *C. pacificum*, both isolated from *Pocillopora* corals like SCF049.01. Nonetheless, its phylogenetic placement is not close enough to *C. proliferum* or *C. vulgare* for it to be placed in either species, suggesting that SCF049.01 belongs to an undescribed species of the C1 radiation within the *Cladocopium* genus.

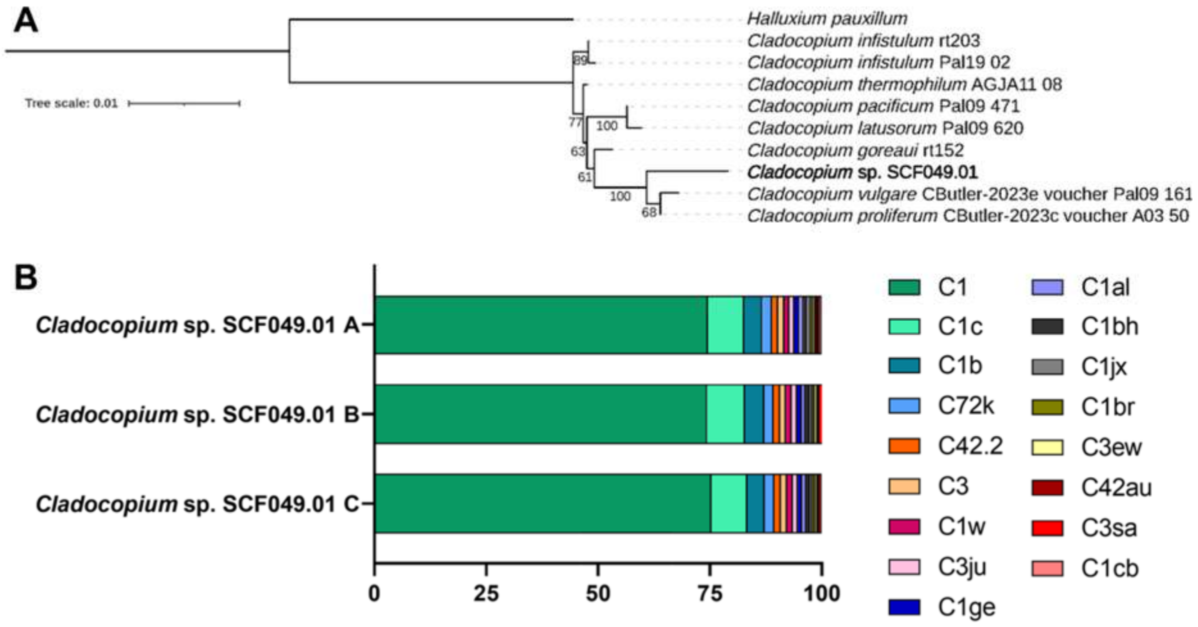


**Figure S1:** Taxonomic placement of *Cladocopium* sp. SCF049.01. **A:** Maximum likelihood phylogeny from aligned concatenated markers (ITS2, LSU, cp23S, *cob*, and *cox1*), showing the relationship of *Cladocopium* sp. SCF049.01 with other described *Cladocopium* species. Bootstrap values (%) based on 1000 replications are provided. **B:** ITS2 metabarcoding showing Symbiodiniaceae ITS2 types of three replicate flasks of *Cladocopium* sp. SCF049.01.

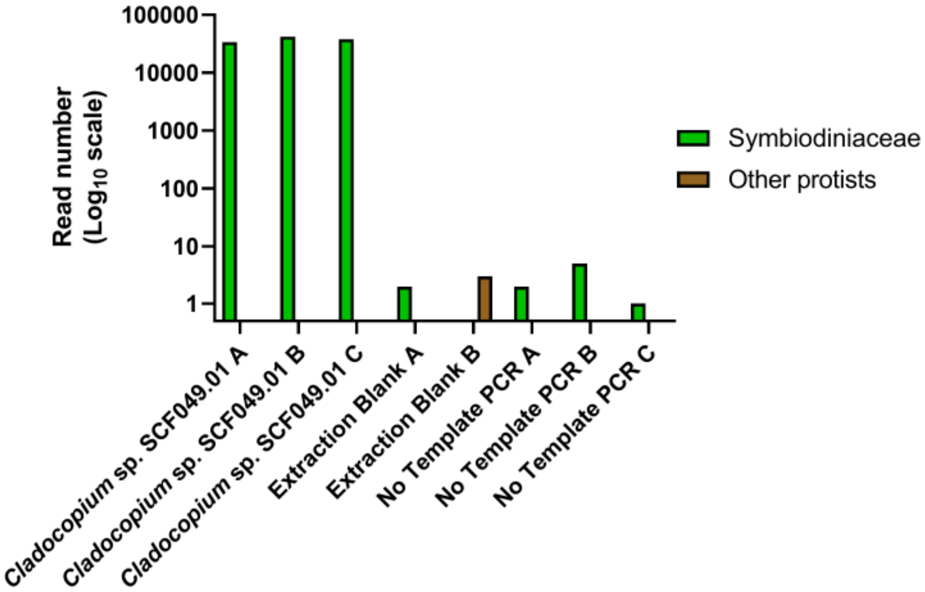


**Figure S2:** 18S rRNA gene metabarcoding showing the protist composition of three replicate flasks of *Cladocopium* sp. SCF049.01.

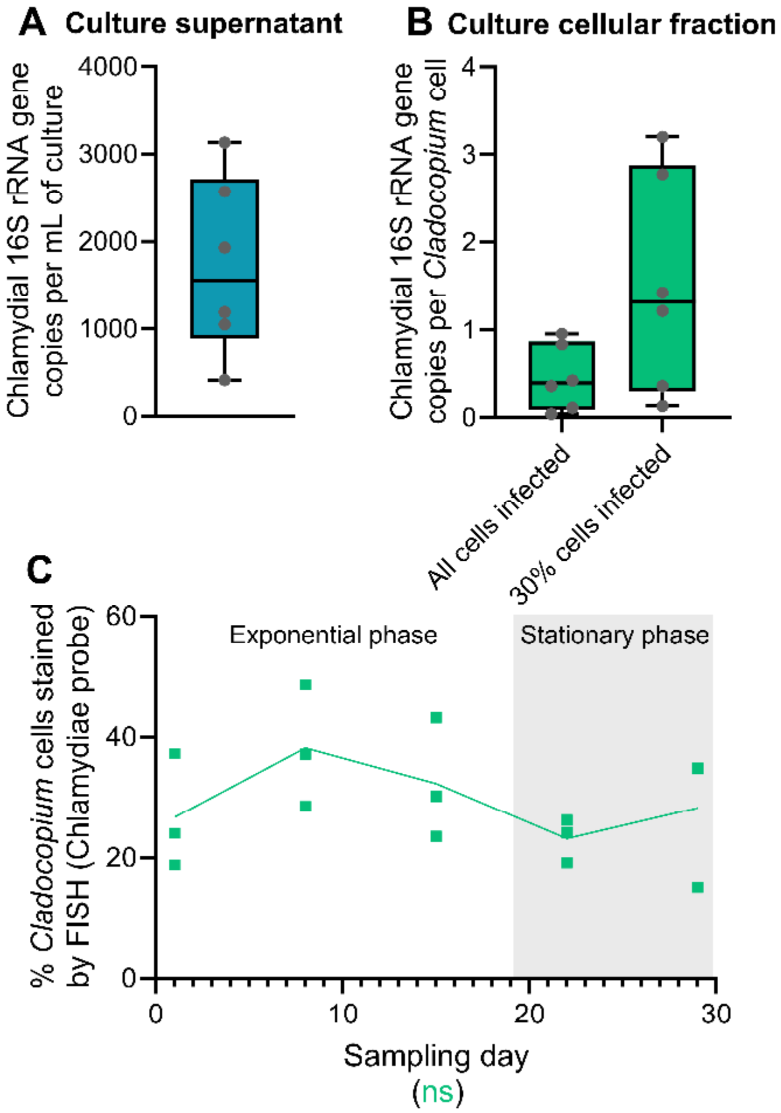


**Figure S3:** Chlamydial infectivity in *Cladocopium* sp. SCF049.01. **A, B:** Chlamydial abundance in culture supernatant (A) or cellular fraction (B), measured by dPCR. For the cellular fraction (B), gene copy data was normalized by *Cladocopium* cell numbers, with the assumption that either all cells or 30% of cells (see C) harbored chlamydiae. Boxes represent first quartile to third quartile for six independent replicates (also shown individually), the middle lines represent the medians, and the whiskers represent the minimum and maximum values. **C:** Proportion of *Cladocopium* sp. cells stained by FISH with the chlamydiae-specific probe Chls523, as analyzed by flow cytometry. Each point represents one of three replicate flasks, with lines connecting the means. Gray-shaded areas represent dark time, while white areas represent light time. The effect of sampling day is indicated under the plot with its corresponding color, based on a Kruskal-Wallis test (ns: $p > 0.05$).

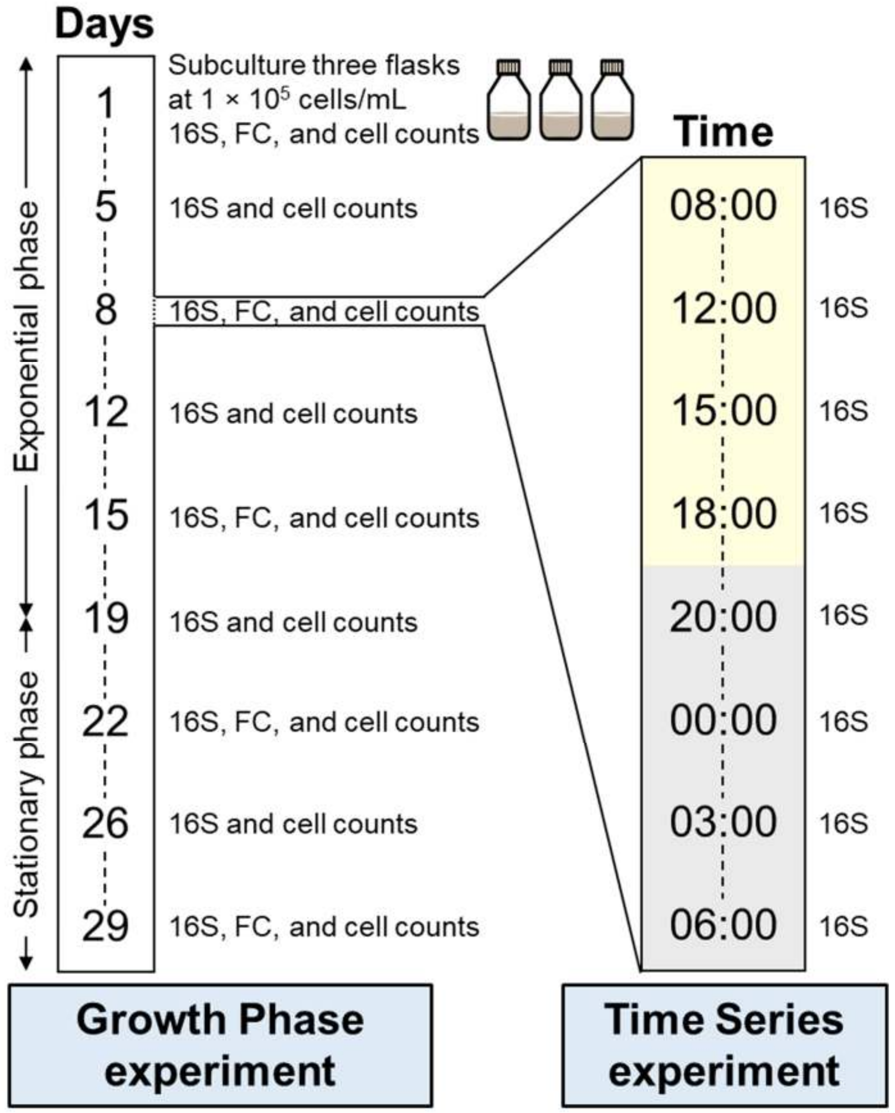


**Figure S4:** Experimental design of the growth phase (left) and time series experiments (right; day 8 of the growth phase experiment). On sampling days or times, flasks were sampled for 16S rRNA gene metabarcoding ('16S'), flow cytometry ('FC', growth phase experiment only) and cell counts (growth phase experiment only). Yellow shading represents daytime sampling and grey shading represents nighttime sampling.

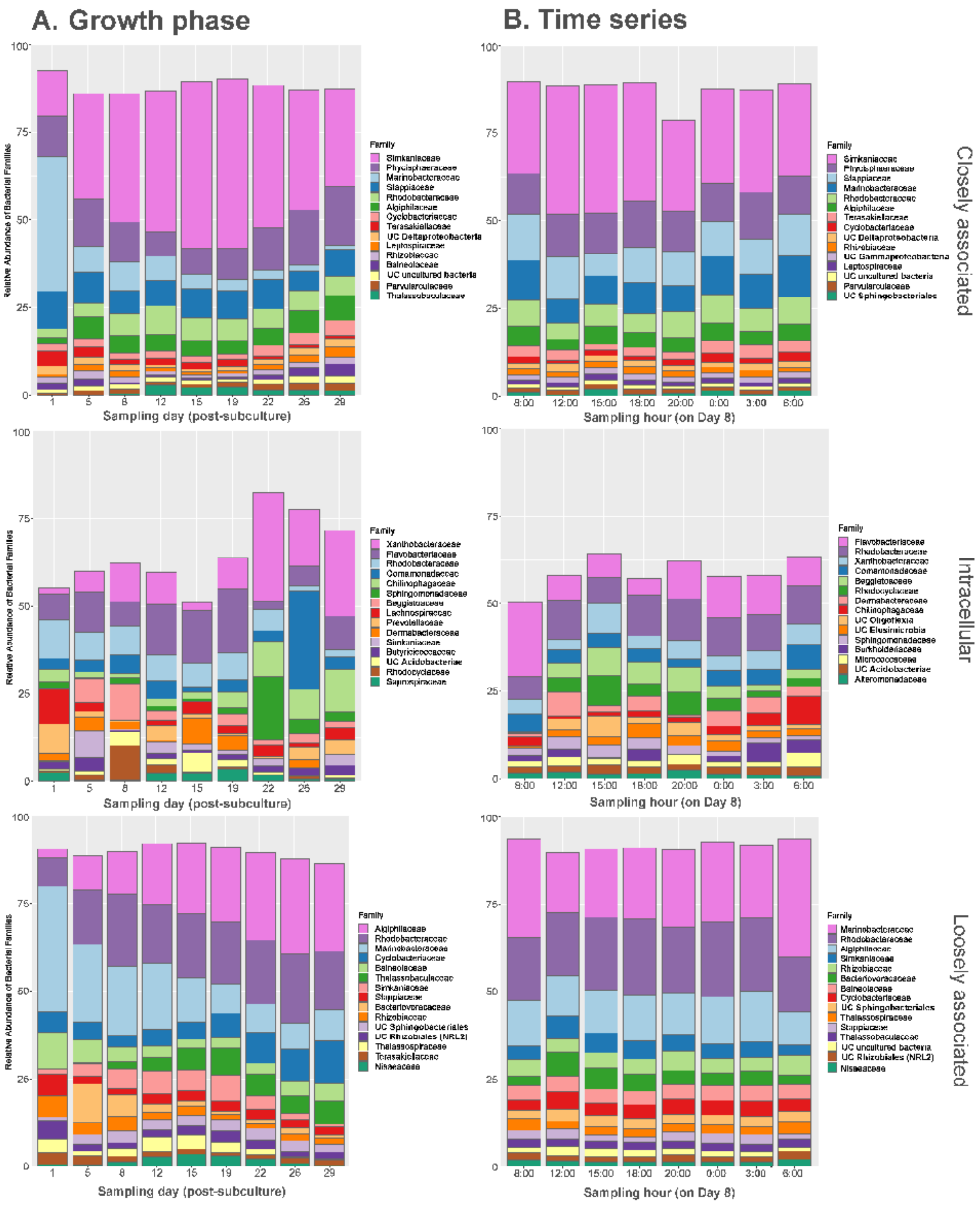


**Figure S5:** Relative abundance of the 15 most abundant bacterial families in *Cladocopium* sp. SCF049.01's closely associated (top), intracellular (middle), and loosely associated (bottom) communities in the growth phase (A) and time series (B) experiments. For each strain sampling time, three independent replicate flasks were merged. UC: unclassified.

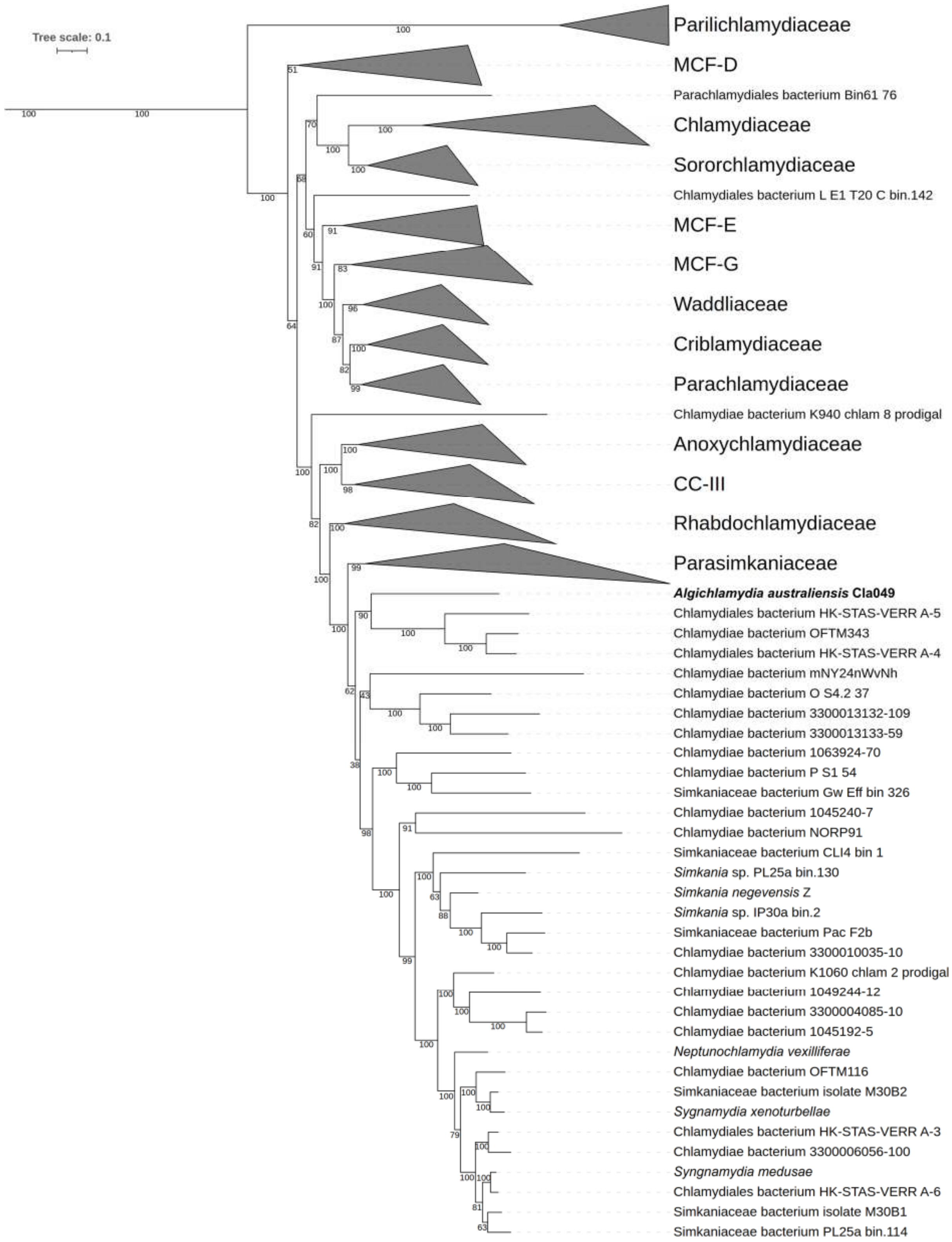


**Figure S6:** Chlamydial maximum likelihood phylogeny based on 15 conserved gene markers (Table S4) in 169 chlamydial genomes (Table S5). Confidence values based on 1000 bootstrap replicates are provided. This tree was calculated under the posterior mean **s**ite frequency (PMSF) model [3] using the tree in Figure 2 as seed. MCF: metagenomic chlamydial family; CC-III: chlamydiae clade III.

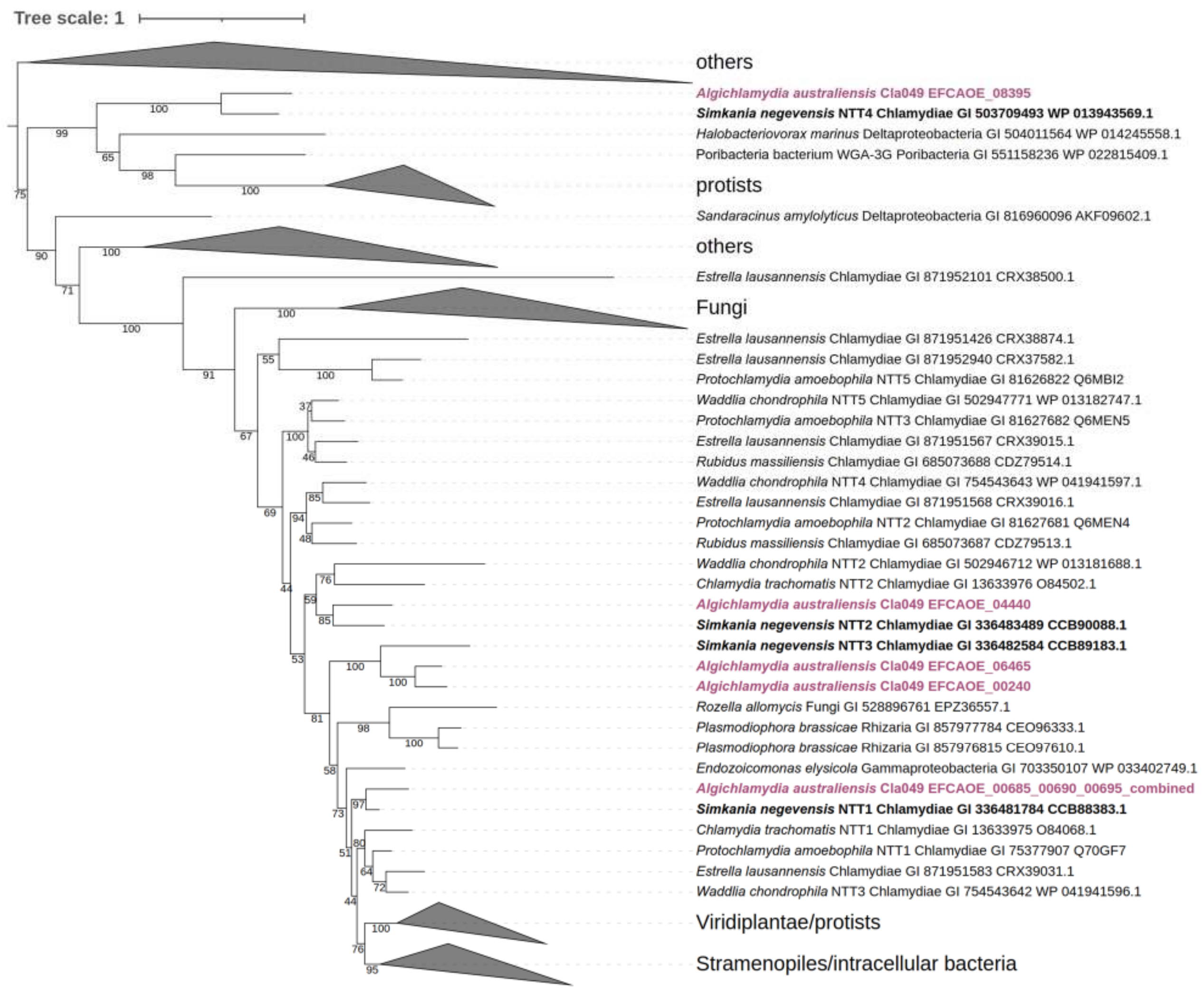


**Figure S7:** Phylogenetic tree of nucleotide transport proteins (NTTs) of *Algichlamydia australiensis* Cla049, chlamydiae, and other organisms [4]. Confidence values based on 1000 bootstrap replicates are provided. The NTT1 homolog in *A. australiensis* is a pseudogene.

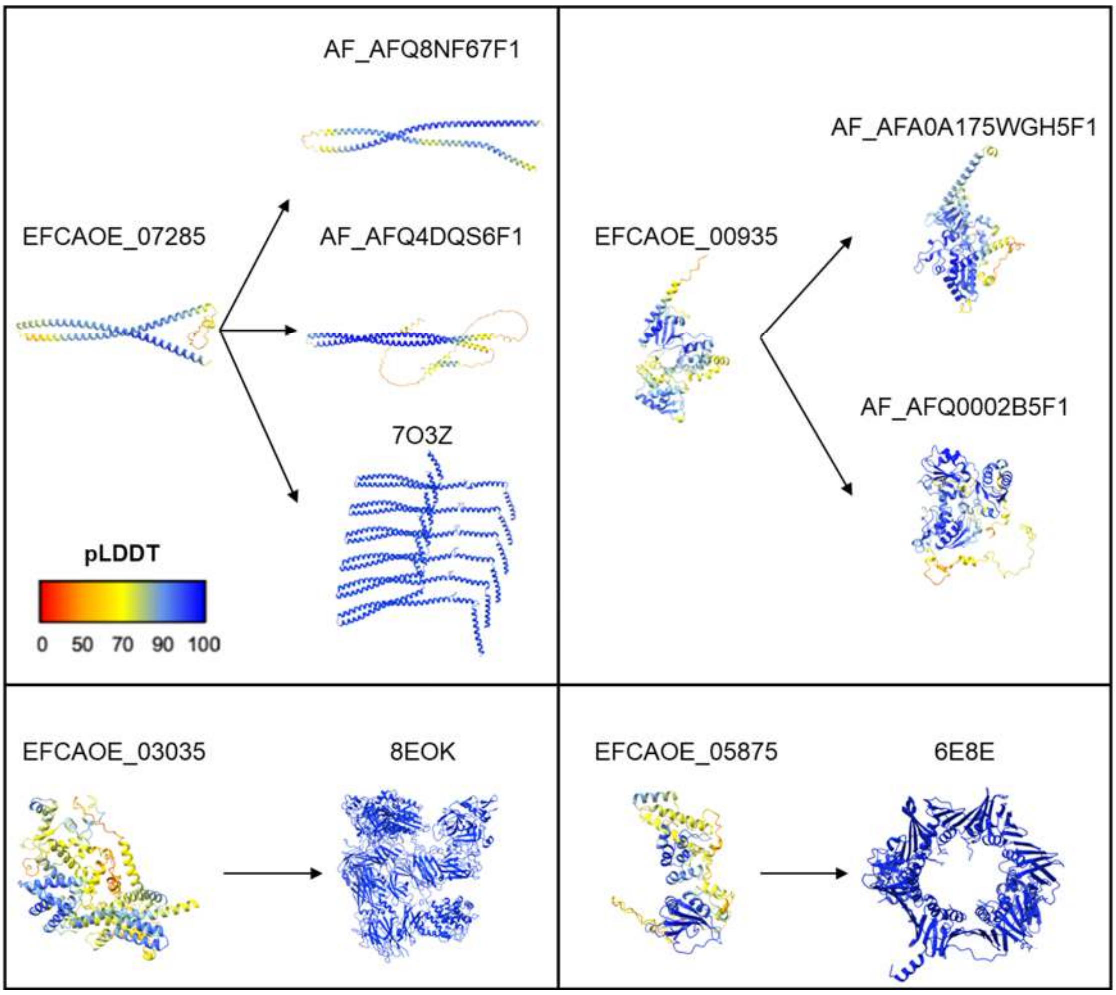


**Figure S8:** Predicted structures of four representative genes of OGs unique to *Algichlamydia australiensis* Cla049. Only structures where global pLDDT > 70 are shown here, along with the closest protein structures. See Table S11 for additional information.

**Table S1:** Sequencing statistics for the two 16S rRNA gene metabarcoding experiments analyzed in this study. The first column ("Growth Phase") refers to Figure 1C and Table S2A, and the second column ("Time Series") refers to Figure 1D Table S2B.

| Experiment | Growth Phase | Time Series |
|---|---|---|
| **Total Samples (negative controls)** | 98 (17) | 89 (17) |
| **Raw reads** | 3897444 | 3676726 |
| **Reads after merging, denoising and chimera filtering** | 3205712 | 2949638 |
| **Contaminating ASVs** | 29 | 33 |
| **Contamination (%)** | 5.99 | 9.88 |
| **Samples kept for analysis** | 76 | 69 |
| **ASVs after decontamination** | 493 | 2474 |
| **Read per sample** | 40985 | 41642 |

**Table S2:** Relative abundance of chlamydial ASVs in *Cladocopium* sp. SCF049.01 samples of the growth phase experiment (A) and the time series experiment (B). This data is summarized in Figure 1C-D.

(attached)

**Table S3:** List of conserved genes on chlamydial plasmids and their locus in the genome of *Algichlamydia australiensis* Cla049.

| Gene | Function | Locus | Contig |
|---|---|---|---|
| SNE_B24960_pgp1 | replicative DNA helicase | EFCAOE_00120, EFCAOE_03395 | **1 (Plasmid)**, 3 |
| SNE_B24950_pgp2 | virulence plasmid protein | EFCAOE_00130 | **1 (Plasmid)** |
| SNE_B24970_pgp5/parA | chromosome partitioning | EFCAOE_03160, EFCAOE_03815 | 3 |
| SNE_B24980_pgp6 | induction of IFN gamma mediated host cell response | EFCAOE_03830 | 3 |

**Table S4:** Marker Non-supervised Orthologous Group (NOG) proteins used for chlamydial phylogenetic analysis.

| **NOG** | **NOG category** | **NOG description** |
|---|---|---|
| COG0064 | J | Aspartyl-tRNA (Asn)/glutamyl-tRNA (Gln) amidotransferase subunit B |
| COG0092 | J | Ribosomal protein S3 |
| COG0233 | J | Ribosome recycling factor |
| COG0290 | J | Translation initiation factor IF-3 |
| COG0292 | J | Ribosomal protein L20 |
| COG0323 | L | DNA mismatch repair protein MutL |
| COG0335 | J | Ribosomal protein L19 |
| COG0342 | U | Preprotein translocase subunit SecD |
| COG0468 | L | DNA recombination/repair protein RecA |
| COG0532 | J | Translation initiation factor IF-2 |
| COG0536 | DL | GTPase Obg involved in cell cycle, chromosome segregation and ribosome assembly |
| COG0706 | M | Membrane protein insertase YidC |
| COG1185 | J | Polyribonucleotide nucleotidyltransferase Pnp |
| COG1530 | J | Ribonuclease G or E |
| COG1663 | M | Tetraacyldisaccharide-1-P 4’-kinase LpxK |

**Table S5:** List of chlamydial genomes used for phylogenetic analyses.

(attached)

**Table S6:** Average amino acid identity (AAI) of the Cla049 MAG with other Simkaniaceae and Parasimkaniaceae genomes. Additional data on the reference genomes is available in Table S5.

(attached)

**Table S7:** Detailed Prokka and eggNOG-mapper annotations for *Algichlamydia australiensis* Cla049.

(attached)

**Table S8:** List of complete and incomplete pathways in *Algichlamydia australiensis* Cla049.

| **Category** | **Pathways > 80% complete** | **Incomplete pathways** |
|---|---|---|
| **Carbon metabolism** | Glycolysis (Embden-Meyerhof pathway) | Entner-Doudoroff pathway |
| | Pentose phosphate pathway | PRPP biosynthesis |
| | Citrate cycle (TCA cycle) | |
| | Pyruvate oxidation | |
| | Glycogen biosynthesis | |
| **Nitrogen metabolism** | | Denitrification |
| | | Dissimilatory nitrate reduction |
| | | Assimilatory nitrate reduction |
| **Nucleotide biosynthesis** | | De novo purine biosynthesis |
| | | De novo pyrimidine biosynthesis |
| **Amino acid biosynthesis** | Glycine-serine interconversion | Asparagine biosynthesis |
| | Aspartate biosynthesis | Glutamine biosynthesis |
| | Alanine biosynthesis | Serine biosynthesis |
| | Glutamate biosynthesis | Threonine biosynthesis |
| | Lysine biosynthesis | Ectoine biosynthesis |
| | Shikimate pathway | Cysteine biosynthesis |
| | | Methionine biosynthesis |
| | | Valine/isoleucine biosynthesis |
| | | Leucine biosynthesis |
| | | Arginine biosynthesis |
| | | Proline biosynthesis |
| | | Histidine biosynthesis |
| | | Tryptophan biosynthesis |
| | | Phenylalanine biosynthesis |
| | | Tyrosine biosynthesis |
| | | Spermidine biosynthesis |
| | | Putrescine biosynthesis |
| | | Glutathione biosynthesis |
| **Cofactor biosynthesis** | Menaquinone biosynthesis | Thiamine biosynthesis |
| | | Riboflavin biosynthesis |
| | | Pyridoxal biosynthesis |
| | | Pantothenate biosynthesis |
| | | Pimeloyl-ACP biosynthesis |
| | | Cobalamin biosynthesis |
| | | Biotin biosynthesis |
| | | Tetrahydrofolate biosynthesis |
| | | Siroheme biosynthesis |
| | | Heme biosynthesis |
| | | NAD biosynthesis |
| | | Coenzyme A biosynthesis |
| | | Ubiquinone biosynthesis |

**Table S9:** List of predicted secondary metabolites in *Algichlamydia australiensis* Cla049.

| Contig name | Type | Closest biosynthetic gene cluster ID | Closest biosynthetic gene cluster compound | Core biosynthetic gene | Product |
|---|---|---|---|---|---|
| contig_2 | NRPS-like | BGC0002711 | nostovalerolactone | EFCAOE_01065 | PlsC domain-containing protein |
| contig_3 | NRPS-like | BGC0002711 | nostovalerolactone | EFCAOE_04950 | hypothetical protein |

**Table S10:** List of hallmark chlamydial genes found in *Algichlamydia australiensis* Cla049, related to virulence processes, and their putative functions in host-chlamydiae interactions.

(attached)

**Table S11:** List of genes present *Algichlamydia australiensis* Cla049 and absent from all other chlamydiae. The closest proteins based on AlphaFold structural predictions are provided. pLDDT: predicted local distance difference test score (high confidence if pLDDT > 70). Structures with low confidence are italicized. Only RCSB hits with pLDDT > 70 were considered.

(attached)

**Table S12:** Relative abundance of *Algichlamydia australiensis* Cla049 in from Symbiodiniaceae (A) and cnidarian (B) 16S rRNA gene metabarcoding samples. The Symbiodiniaceae data was obtained from a previous study that analyzed bacterial community composition in 11 Symbiodiniaceae cultures [5]. The cnidarian data was obtained from a recent dataset combining 186 cnidarian microbiome studies and 12,009 cnidarian samples [6].

(attached)

**Table S13:** List of contaminants identified in the 16S rRNA gene metabarcoding data, and their abundance in *Cladocopium* sp. SCF049.01 samples, in the growth phase experiment (A) and the time series experiment (B).

(attached)
